# piRNA direct the chromatin reader to its genomic targets

**DOI:** 10.64898/2026.05.08.723782

**Authors:** Yicheng Luo, Wenjiang Zhou, Zhaohui Jin, Baira Godneeva, Xiawei Huang, Haoqing Wang, Peng He, Ying Huang, Alexei A. Aravin

**Author notes:** To whom correspondence should be addressed: Yicheng Luo Alexei A. Aravin.

## Abstract

The chromatin reader Rhino (Rhi) is an HP1-paralog and master regulator of piRNA biogenesis in *Drosophila* that drives the transcription of piRNA precursors. While Rhi binds the heterochromatic mark H3K9me3, this interaction is insufficient to explain its specific occupancy, as Rhi is excluded from the majority of H3K9me3-enriched loci. Instead, we find that Rhi recruitment is orchestrated by several competing pathways, including the piRNA-guided protein Panoramix (Panx) and the DNA-binding protein Kipferl (Kipf). We demonstrate that Panx functions as a piRNA-guided modular scaffold that directs Rhi to its targets through a dual-mode mechanism: its N-terminus recruits the H3K9 methyltransferase machinery, while its C-terminus directly binds the Rhi chromodomain. While H3K9me3 deposition provides a necessary foundation, it acts in synergy with the direct Panx-Rhi physical bridge to drive robust Rhi occupancy. Furthermore, Rhi recruitment is amplified by the cytoplasmic inheritance of maternal piRNAs, forming a transgenerational feed-forward loop. Our results reveal that the piRNA pathway imparts sequence-specificity to the H3K9me3 code, utilizing a direct physical bridge to program the genomic distribution of the chromatin reader Rhi.

## Introduction

The regulation of eukaryotic genomes relies heavily on the “histone code,” a complex network of post-translational modifications that demarcate distinct chromatin domains. These marks act as docking sites for effector proteins, or “chromatin readers,” which translate the chromatin landscape into functional biological outcomes. For example, H3K9me2/3 marks define heterochromatin domains associated with transcriptional repression, a function mediated by HP1 family proteins that bind this mark through their chromodomain^1–6^. However, the classical lock-and-key model—where a specific histone mark perfectly predicts the binding of its cognate reader—frequently fails to explain the precision of chromatin regulation *in vivo*^6,7^. Indeed, genomic profiling frequently reveals a striking discordance: readers are often excluded from regions where their target marks are abundant, or occupy sites where the marks are depleted^7–10^. This raises a fundamental question: how does a chromatin reader achieve locus-specific targeting?

This paradox is strikingly illustrated in the *Drosophila* germline by the Heterochromatin Protein 1 (HP1) family of chromatin readers. Canonical HP1a utilizes its chromodomain to bind H3K9me3, occupying H3K9me3-rich heterochromatin to maintain transcriptional repression^1–4,10^. In stark contrast, its germline-specific paralog, Rhino (Rhi), binds the exact same H3K9me3 mark^11,12^; however, its occupancy is restricted to a subset of loci that encode piRNAs^5,7,10^—short, non-coding RNAs that act as guides for the repression of transposable elements (TEs) to protect the genome against their activity. On chromatin, Rhi interacts with several partners to drive the non-canonical transcription and nuclear export of piRNA precursors^7,13–19^. Because Rhi binding functionally defines piRNA-generating loci, its precise localization is essential. If Rhi were mislocalized, it could trigger the aberrant piRNA targeting of protein-coding genes, while a failure to bind target loci would unleash TE activity and cause severe genome damage^5,7,10^.

In the nucleus, the piRNA pathway directs the co-transcriptional silencing when piRNA-loaded Piwi complexes recognize nascent TE transcripts via RNA-RNA complementarity^20–22^. This piRNA-guided repression depends on the Piwi partner Panoramix (Panx), which recruits the histone methyltransferase SetDB1 (Eggless) in a SUMO-dependent manner to deposit H3K9me3 on the target loci^23–31^. Thus, piRNAs direct the *de novo* establishment of the H3K9me3 mark, which in turn was presumed to provide the primary anchor for Rhi binding and the generation of new piRNAs.

As Rhi sits at the top of the hierarchy that defines piRNA loci, understanding the molecular mechanisms governing its chromatin localization is crucial. Recent studies identified the DNA-binding zinc finger protein Kipferl (Kipf) as a factor that recruits Rhi to guanosine-rich motifs^10^. Another study implicated a different histone mark, H3K27me3, in recruiting Rhi to chromatin^32^. How these mechanisms interact with the H3K9me3-binding activity of the Rhi chromodomain, and whether they are sufficient to explain Rhi targeting *in vivo*, remained unclear. Here, we resolve the paradox of Rhi specificity by demonstrating that its recruitment is not an intrinsic property of the histone code, but is directly guided by the piRNA pathway. We find that the Piwi partner Panx acts as a modular, two-pronged scaffold: its N-terminus recruits the H3K9 methyltransferase machinery, whereas its structured C-terminal region binds Rhi directly. Although the resulting H3K9me3 mark provides a necessary foundation, it is this Panx-Rhi physical bridge that ensures specific recruitment. Finally, we demonstrate that Panx tethering is sufficient to trigger *de novo* piRNA biogenesis at a target locus, a state which is subsequently maintained across generations by maternally inherited piRNAs. Together, our findings highlight Panx as a key link connecting transcriptional repression to piRNA biogenesis, revealing how small RNAs superimpose sequence specificity onto the broader histone code to program the exact genomic distribution of a chromatin reader.

## Results

### H3K9me3 recognition is insufficient to explain the specific genomic recruitment of Rhi

The paralogous proteins HP1 and Rhi share a common domain architecture. The chromodomains of both HP1 and Rhi bind H3K9me3 *in vitro*^11,12^ and are required for their chromatin localization *in vivo*^5,7^. These findings suggest that recognition of the H3K9me3 mark guides both proteins to their respective chromatin targets. However, despite this shared recruitment mechanism, previous studies have reported that these two proteins possess distinct genomic binding profiles^10^. Because HP1 is ubiquitously expressed while Rhi is restricted to germ cells, it remains unclear whether these reported differences reflect intrinsic binding preferences or are merely a consequence of their divergent expression across different cell types. To resolve this, we utilized immunofluorescence analysis in nurse cell nuclei—where both proteins are naturally co-expressed—to compare their localization within a shared chromatin environment. Immunofluorescence analysis revealed that while both proteins localize to nuclear foci that overlap with H3K9me3-rich heterochromatin, their distributions only partially coincide (Fig. 1A). Specifically, HP1 is present in all H3K9me3 foci (Pearson’s *r* = 0.873), whereas Rhi is restricted to a specific subset of these foci (Pearson’s *r* = 0.567).

**Figure 1.**
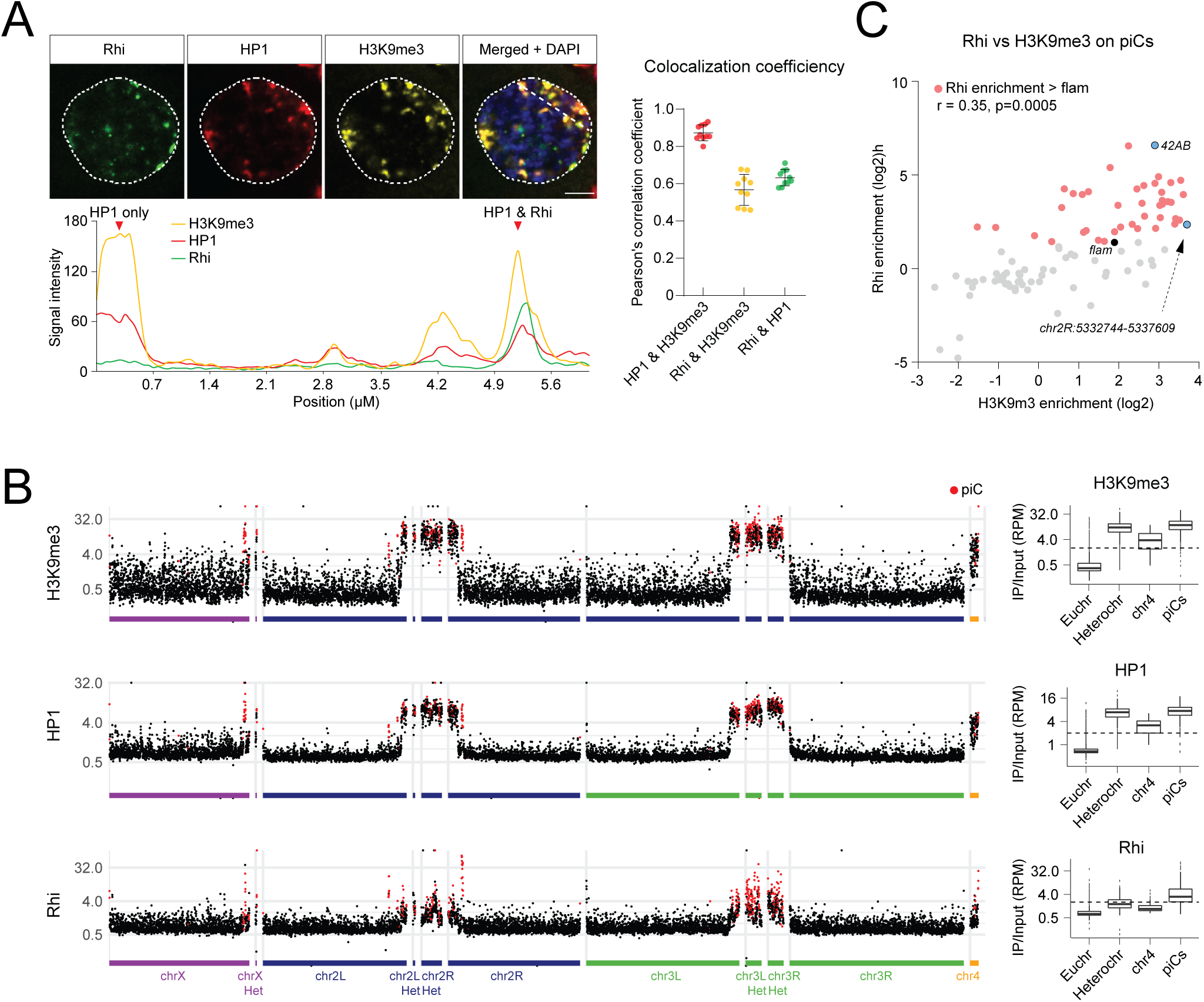
Rhi and HP1 have distinct localization patterns within H3K9me3-rich heterochromatin. **(A) Rhi, HP1, and H3K9me3 localize to partially overlapping foci in nurse cell nuclei.** GFP-Rhi, HP1, and H3K9me3 were visualized by immunofluorescence in nurse cell nuclei. Scale bar represents 2 µm. The plot profile shows fluorescence intensity along the marked section, with foci containing only HP1, or both HP1 and Rhi, indicated. Pearson’s correlation coefficients (*r*) quantify the co-localization between HP1 and H3K9me3 (red), Rhi and H3K9me3 (yellow), and Rhi and HP1 (green) (*n* = 10 nuclei). **(B) Genome-wide profiling reveals distinct binding patterns of Rhi and HP1.** ChIP-seq profiles of H3K9me3, HP1a, and Rhi across *D. melanogaster* chromosomes. Data points correspond to input-normalized ChIP signals in 10 kb genomic intervals. Intervals overlapping piCs are marked in red. Boxplots show the distributions of ChIP signals measured in 10 kb genomic intervals within euchromatin (Euchr), pericentric heterochromatin (Heterochr), chromosome 4 (chr4) and piRNA clusters (piCs). Results were averaged from two biological replicates. **(C) Rhi enrichment at piCs shows a weak correlation with H3K9me3 levels.** The scatter plot shows the enrichment of H3K9me3 and Rhi across piRNA clusters, with Rhi-enriched clusters highlighted in red and representative piCs labeled in blue. ChIP signals were input-normalized, and results were averaged from two biological replicates. Pearson’s correlation coefficient (*r* = 0.35) and *p*-value (*p* = 0.0005) indicate a weak correlation between H3K9me3 and Rhi signals.

To further characterize the genomic distribution of HP1 and Rhi, we performed parallel ChIP-seq profiling using *Drosophila* ovary chromatin. We conducted these experiments using both antibodies against the native proteins and transgenic flies expressing GFP-tagged HP1 and Rhi under the control of a germline-specific promoter. The latter approach allowed us to profile the chromatin occupancy of both proteins within the same cell type and under identical conditions, utilizing the same anti-GFP antibody to eliminate potential bias. Consistent across all experimental setups, genome-wide profiling corroborated our immunofluorescence results: HP1 occupancy strongly correlated with the H3K9me3 mark across the entire genome. In contrast, Rhi was restricted to a specific subset of H3K9me3-enriched loci, which also exhibited high levels of HP1 (Fig. 1B and S1A,B). Combined, IF and ChIP-seq show a drastic difference in chromatin localization of HP1 and Rhi. HP1 occupancy closely recapitulates the H3K9me3 landscape and therefore can be guided by binding to this mark. In contrast, Rhi localization is more specific indicating that binding of H3K9me3 mark is necessary but not sufficient to explain Rhi binding to chromatin.

Although Rhi is excluded from many H3K9me3-enriched regions, we sought to determine whether Rhi occupancy correlates with H3K9me3 density within the specific domains where it is bound. In agreement with previous reports^5,7^, our genome-wide profiling confirmed that Rhi is localized to dual-strand piRNA clusters (piCs) (Fig. 1B). However, we observed a weak correlation between the levels of Rhi and H3K9me3 across 41 Rhi-enriched piCs (Pearson’s *r* = 0.35; Fig. 1C). For instance, the cluster exhibiting the highest level of H3K9me3 (chr2R: 53,327,444–53,337,609) showed low Rhi enrichment, whereas the cluster with the highest Rhi occupancy, *42AB*, possessed only intermediate levels of H3K9me3. Thus, even at genomic loci enriched for Rhi, its abundance does not strictly track with H3K9me3 density. These results further underscore the existence of additional, H3K9me3-independent mechanisms that guide the precise recruitment of Rhi to its chromatin targets.

### Panx directs Rhi to a distinct set of genomic targets in competition with Kipf and E(z)

What mechanisms underlie the selective recruitment of Rhi to specific H3K9me3-enriched domains? Recent studies have identified Kipf, a ZAD zinc-finger protein, as a Rhi-interacting factor that facilitates its chromatin recruitment^10^. It has also been proposed that the co-occurrence of H3K9me3 and H3K27me3 histone marks may guide Rhi to specific piRNA clusters, although the precise mechanism remains to be elucidated^32^. Alternatively, the tight link between Rhi occupancy and piRNA biogenesis suggests a feed-forward model: piRNAs, by recognizing complementary nascent transcripts, may direct the deposition of Rhi to the genomic loci from which they originate^7,33^. Indeed, piRNAs guide the nuclear Piwi protein to establish H3K9me3 marks that provide a platform for Rhi binding^7^. This process requires Panx, which interacts directly with Piwi and is essential for piRNA-directed H3K9me3 establishment^23,24^. Consequently, if the piRNA pathway actively directs Rhi to chromatin, Rhi occupancy should be disrupted upon Panx depletion.

To evaluate the requirement of Panx for Rhi recruitment, we performed genome-wide ChIP-seq analysis of Rhi occupancy in ovaries following *Panx* germline knockdown (*Panx* GLKD). We first assessed changes in Rhi enrichment across various TE families, which revealed a significant reduction in Rhi occupancy at several TEs upon Panx depletion (Fig. 2A). The most pronounced loss of Rhi was observed for the telomere-associated TEs *HeT-A* (17.5-fold), *TART-A* (15.6-fold), and *TAHRE* (4.5-fold). In addition to TEs, we examined Rhi levels at piCs. Many dual-strand piCs exhibited a marked loss of Rhi following *Panx* GLKD. Specifically, 14 of the 41 (34%) Rhi-enriched clusters displayed both a reduction in Rhi occupancy and a greater than 2-fold decrease in piRNA abundance, suggesting that the loss of Rhi at these loci disrupts piRNA biogenesis (Fig. 2B and 2C). Notably, the changes in Rhi levels and piRNA production were correlated; no decrease in piRNA levels was detected at piCs where Rhi occupancy was maintained despite Panx depletion. Collectively, these results indicate that Panx is essential for both Rhi deposition and subsequent piRNA biogenesis at a specific subset of genomic loci.

**Figure 2.**
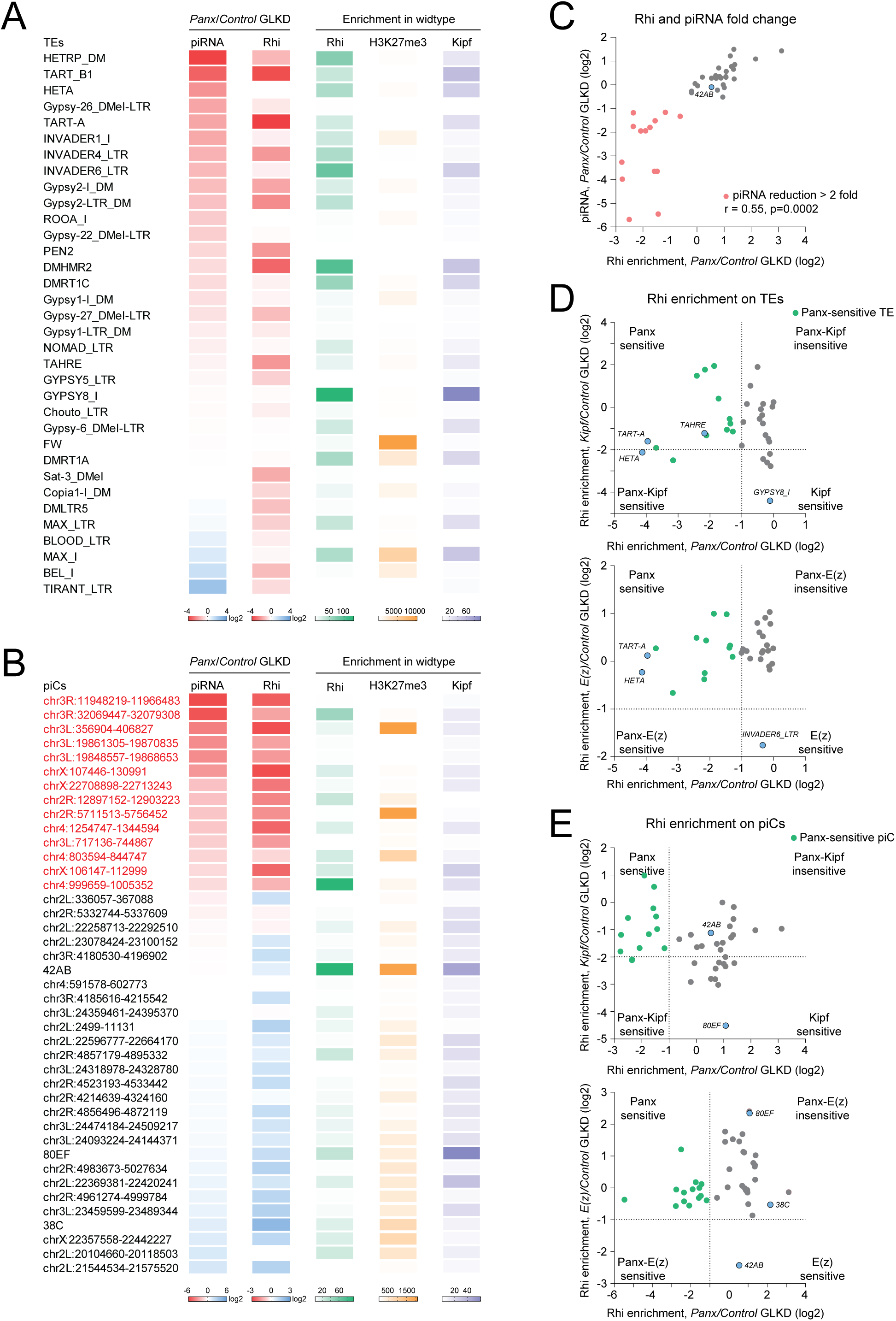
Panx is required for Rhi localization on TE and piRNA clusters in competition with Kipf and E(z) (A) Panx is required for Rhi occupancy at a subset of TE families. The heatmap shows the enrichment of Rhi, Kipf, and H3K27me3 signals across different TE families, as well as changes in Rhi occupancy and piRNA levels upon *Panx* GLKD. Only TE families with a Rhi signal that exceeds that of the Flam cluster and decreases upon *Panx* GLKD are shown. The results were averaged from two biological replicates. **(B) Panx promotes Rhi occupancy and piRNA production at a subset of piCs.** The heatmap shows the enrichment of Rhi, Kipf, and H3K27me3 signals across different piCs, as well as changes in Rhi occupancy and piRNA levels upon *Panx* GLKD. Only piCs with a Rhi signal exceeding that of the *Flam* cluster are shown. The results were averaged from two biological replicates. **(C) Loss of Rhi upon Panx depletion correlates with reduced piRNA output**. The scatter plot shows changes in Rhi enrichment and piRNA levels at Rhi-enriched piCs upon *Panx* GLKD. Clusters with a >2-fold reduction in piRNA levels are highlighted in red. piRNA levels are normalized to *Flam* piRNAs. The results were averaged from two biological replicates. Pearson’s correlation coefficient (*r*) and *p*-value (*p* = 0.0002) are indicated. **(D) Panx, Kipf, and E(z) have distinct effects on Rhi levels at TEs.** The scatter plots compare fold changes in Rhi occupancy across TE families upon Kipf, E(z), and Panx depletions. *Panx*-sensitive TEs (green dots) are defined by a >2-fold loss of Rhi upon *Panx* GLKD. *Kipf*- and *E(z)*-sensitive TEs are defined by a >4-fold and >2-fold loss of Rhi upon their respective knockdowns. Results were averaged from two biological replicates. Representative TE families are indicated. **(E) Panx, Kipf, and E(z) have distinct and frequently opposite effects on Rhi levels at piCs.** The scatter plots compare fold changes in Rhi occupancy across Rhi-enriched piCs upon Kipf, E(z), and Panx depletions. *Panx*-sensitive piCs (green dots) are defined by a >2-fold loss of Rhi upon *Panx* GLKD. *Kipf*- and *E(z)*-sensitive piCs are defined by a >4-fold and >2-fold loss of Rhi upon their respective knockdowns. Results were averaged from two biological replicates. Representative piCs are indicated.

Although Panx depletion disrupted Rhi occupancy at many genomic loci, the effect was not universal. Certain regions exhibited no significant change or even an increase in Rhi levels, suggesting the involvement of alternative recruitment mechanisms (Fig. 2C). To investigate how the Panx-dependent pathway interacts with other proposed mechanisms—namely, recruitment via the DNA-binding protein Kipferl (Kipf) and the H3K27me3 mark—we integrated our results with published datasets^10,32^ to analyze the occupancy of Kipf and the distribution of H3K27me3. Rhi-enriched TEs and piCs showed high variability in both Kipf levels and H3K27me3 density (Fig. 2A and 2B); notably, we found no correlation between a region’s sensitivity to *Panx* GLKD and its baseline occupancy by Kipf or H3K27me3 (Fig. S2A).

We next compared the regional sensitivity of Rhi to the depletion of Panx, Kipf, and the H3K27 methyltransferase Enhancer of zeste (E(z)). Unlike Panx, which had the most pronounced effect on the telomere-associated TEs *HeT-A*, *TART-A*, and *TAHRE*, depletions of Kipf and E(z) most strongly impacted *Gypsy8* and *Invader6*, respectively (21.3-fold and 3.34-fold reductions in the respective GLKDs). Importantly, Rhi occupancy at both *Gypsy8* and *Invader6* was not significantly affected by Panx depletion (Fig. 2A and 2D). Furthermore, we observed a negative correlation between the sensitivity of piCs to Panx versus Kipf (Fig. 2E). Of the 12 piCs that exhibited a strong (>4-fold) loss of Rhi upon *Kipf* GLKD (“Kipf-sensitive” piCs), only one was sensitive to Panx. Conversely, Panx-sensitive regions were largely insensitive to Kipf loss, indicating that Kipf and Panx regulate distinct sets of piCs. Beyond regulating different loci, Panx and Kipf often exerted opposite effects on the same clusters: *Panx* GLKD triggered an increase in Rhi levels at 9 of the 12 Kipf-sensitive regions. For instance, the *80EF* cluster—which showed a dramatic 23-fold loss of Rhi upon Kipf depletion—exhibited a 2.1-fold increase in Rhi following *Panx* GLKD (Fig. 2E). Collectively, these results suggest that Panx and Kipf participate in two competing recruitment mechanisms; consequently, the disruption of Panx-mediated targeting may lead to the redistribution of Rhi to Kipf-dependent clusters such as *80EF*.

Certain genomic regions, such as the major piC *42AB*, were insensitive to the depletion of both Panx and Kipf. In contrast, Rhi occupancy at *42AB* was reduced following the depletion of E(z), the H3K27 methyltransferase responsible for H3K27me3 deposition. Remarkably, *42AB* was the only region to exhibit a greater than 2-fold decrease in Rhi levels upon *E(z)* GLKD, although 10 of the 14 Panx-sensitive piCs showed modest (<2-fold) reductions (Fig. 2E). Interestingly, while the endogenous *42AB* cluster remained Panx-insensitive, analysis of fly strains harboring unique sequence insertions within *42AB* revealed that Rhi occupancy and H3K9me3 levels at these specific insertion sites decreased upon *Panx* GLKD (Fig. S2B and S2C) followed by a reduction in piRNA biogenesis (Fig. S2D). This observation suggests that the sensitivity of Rhi recruitment to Panx can vary locally, even within the confines of a single piC. Collectively, our results demonstrate that Panx, Kipf, and E(z) are required for Rhi deposition at largely distinct genomic loci. Furthermore, the finding that depletion of these factors often exerts opposing effects on the same regions indicates that they represent competing, rather than cooperative, recruitment mechanisms.

### Panx-mediated Rhi recruitment is partially uncoupled from the H3K9me3 histone mark

How do piRNAs and Panx facilitate the recruitment of Rhi to its chromatin targets? Panx is essential for H3K9me3 deposition downstream of Piwi/piRNA complexes, and H3K9me3, in turn, provides a binding platform for the Rhi chromodomain^11,12,23^. While this suggests that the piRNA pathway recruits Rhi primarily by establishing the H3K9me3 anchor, it does not preclude the possibility of a more direct recruitment mechanism. To explore this, we utilized genome-wide ChIP-seq to analyze alterations in the H3K9me3 mark across piCs and TEs following *Panx* GLKD. Among TE families exhibiting a substantial reduction in Rhi upon Panx depletion, *TART-B1* showed only a minimal change in H3K9me3 levels, whereas *HeT-A* demonstrated a robust reduction (Fig. 3A). Overall, of the seven TE families that lost both Rhi and piRNAs upon *Panx* GLKD, five exhibited no significant loss (<2-fold) of H3K9me3, suggesting that Panx can regulate Rhi occupancy through an H3K9me3-independent mechanism (Fig. 3B).

**Figure 3.**
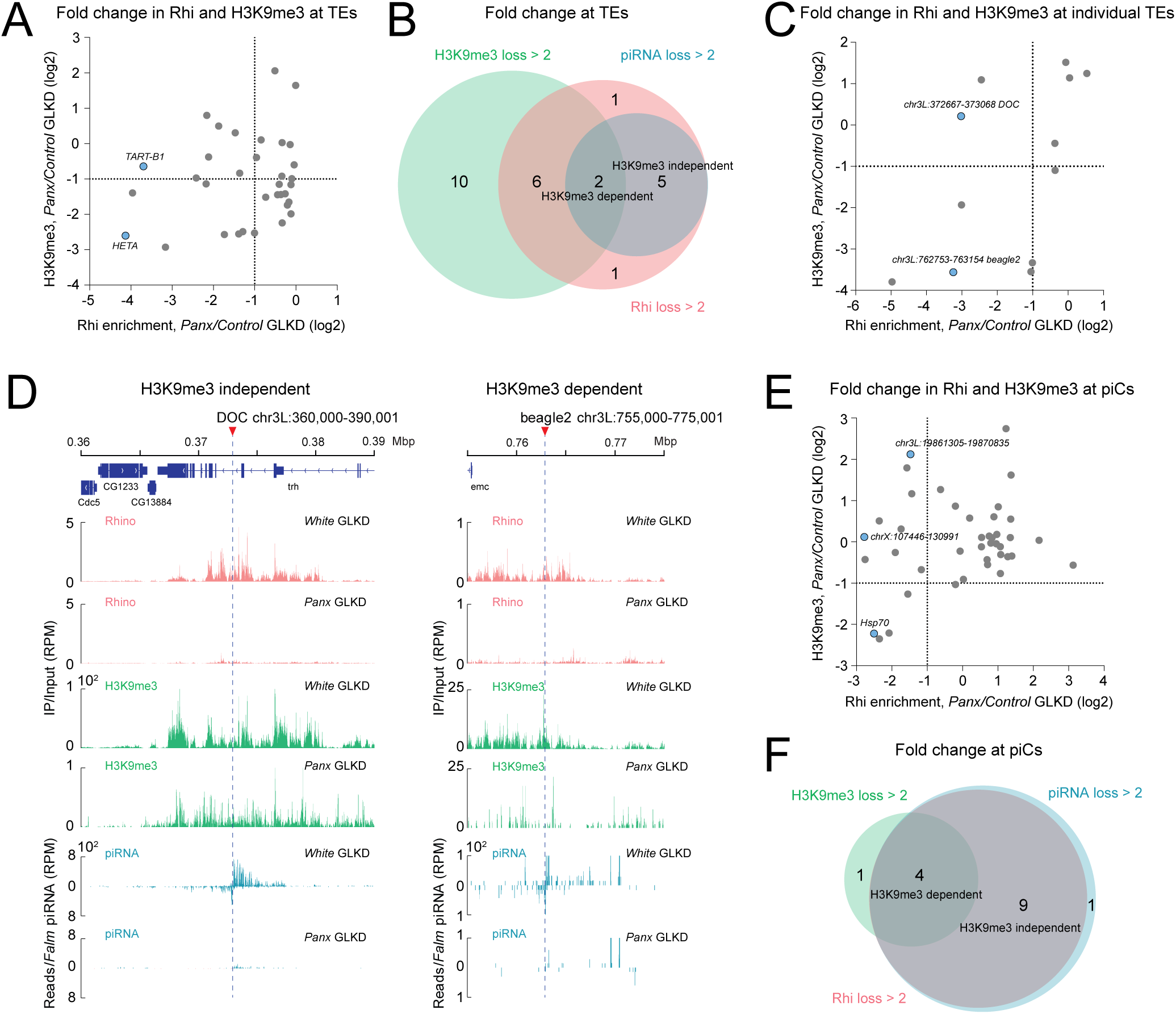
The effect of *Panx* on Rhi recruitment is partially uncoupled from its effect on the H3K9me3 mark. (A) Effect of Panx depletion on Rhi and H3K9me3 levels at TE families. The scatter plot shows fold changes in Rhi occupancy and H3K9me3 levels across TE families upon *Panx* GLKD. Results were averaged from two biological replicates. **(B) Discordant loss of Rhi and H3K9me3 at Panx-sensitive TEs.** The Venn diagram illustrates the number of TE families that exhibit >2-fold reductions in piRNA levels, Rhi occupancy, and H3K9me3 levels upon Panx depletion. Many TE families show a loss of H3K9me3 without an accompanying loss of Rhi, while others show a loss of Rhi without an accompanying reduction in the H3K9me3 mark. **(C) The effect of Panx depletion on individual TE insertions.** The scatter plot shows changes in Rhi occupancy and H3K9me3 levels across the 400 bp flanking regions of individual Rhi-enriched euchromatic TE insertions upon *Panx* GLKD. Results were averaged from two biological replicates. **(D) Examples of the effect of Panx depletion on individual TE insertions.** Shown are profiles of Rhi, H3K9me3, and piRNA levels across the flanking sequences of two euchromatic TE insertions upon *Panx* GLKD. The *DOC* insertion shows a dramatic reduction of both Rhi and piRNA without a loss of the H3K9me3 mark (Left). In contrast, the *beagle2* insertion shows a concomitant loss of Rhi, piRNAs, and H3K9me3 (Right). piRNA levels were normalized to *Flam* piRNAs. **(E) The effect of Panx depletion on Rhi and H3K9me3 levels at piCs.** The scatter plot shows changes in Rhi and H3K9me3 levels at Rhi-enriched piCs upon *Panx* GLKD. Results were averaged from two biological replicates. **(F) The majority of *Panx*-sensitive piCs lose Rhi without a loss of H3K9me3.** The Venn diagram illustrates the number of piCs that exhibit >2-fold reductions in Rhi occupancy, H3K9me3, and piRNA levels upon Panx depletion.

We further examined individual TE insertions within euchromatic regions. Of the 287 euchromatic insertions identified in *Panx* and control (*white*) GLKD strains, 12 exhibited Rhi enrichment and piRNA production in their flanking genomic regions. While these insertions lost Rhi upon Panx depletion, this loss was not consistently associated with a decrease in the H3K9me3 mark (Fig. 3C). For example, at a *DOC* insertion (chr3L: 360,000–390,001), H3K9me3 levels remained stable despite a marked decrease in Rhi binding and piRNA levels (Fig. 3D). In contrast, other loci, such as a *beagle2* insertion (chr3L: 755,000–775,001), showed a strong reduction in H3K9me3 following *Panx* GLKD (Fig. 3D). Collectively, these results demonstrate that Panx is required for Rhi recruitment at stand-alone TE loci via both H3K9me3-dependent and -independent pathways.

Analysis of H3K9me3 levels at piCs upon *Panx* GLKD indicated that a subset of clusters exhibiting Rhi loss also displayed a concomitant decrease in the H3K9me3 mark. For example, the piC at the *hsp70* locus showed a 5.7-fold reduction in H3K9me3 (Fig. 3E), and a similar decrease was observed at transgene insertions within the *42AB* cluster (Fig. S2). However, other clusters exhibited a substantial loss of Rhi without a corresponding reduction in H3K9me3. Notably, the cluster with the most significant Rhi depletion (chrX: 107,446–130,991) showed no reduction in H3K9me3 levels (Fig. 3E). Overall, of the 14 Panx-sensitive piCs, only four displayed a greater than 2-fold reduction in H3K9me3; the majority (64%) experienced Rhi loss without any detectable reduction in the histone mark (Fig. 3E and 3F). Furthermore, three piCs exhibited Rhi loss accompanied by a paradoxical increase in H3K9me3 levels. For instance, one cluster (chr3L: 19,861,305–19,870,835) showed a 2.7-fold reduction in Rhi occupancy and a 10-fold decrease in piRNA levels, despite a 4.5-fold increase in H3K9me3. Conversely, several clusters exhibited a significant reduction in H3K9me3 without a loss of Rhi. Collectively, our analysis of TE families, individual insertions, and piCs demonstrates that Panx is required for Rhi recruitment to a broad range of genomic regions in an H3K9me3-independent manner.

### Panx-mediated Rhi recruitment sustains piRNA production through cytoplasmic inheritance

To determine whether Panx is sufficient for Rhi recruitment, we employed a tethering assay to recruit Panx to a nascent RNA in *Drosophila* germline (Fig. 4A). Tethering of Panx yielded a 5.7- to 6.5-fold enrichment of Rhi, demonstrating that Panx can actively recruit Rhi to chromatin (Fig. 4B). Consistent with previous reports^23,24^, Panx recruitment also led to the enrichment of H3K9me3 and HP1 (Fig. 4B) and triggered transcriptional silencing of the reporter (Fig. 4C). Thus, the presence of Panx is sufficient to recruit both HP1 and Rhi and is associated with the *de novo* deposition of the H3K9me3 mark.

**Figure 4.**
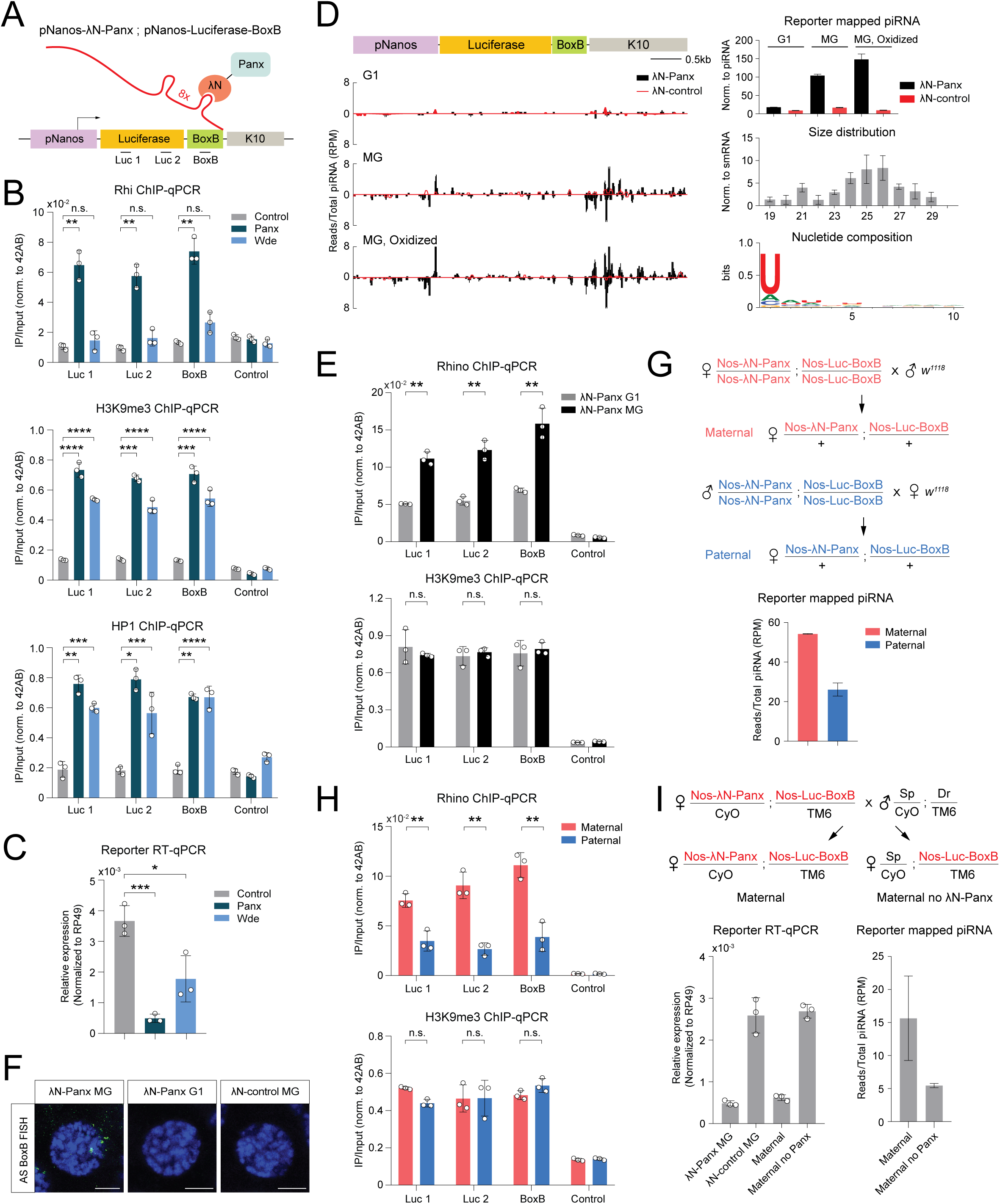
Tethering of Panx to a reporter triggers Rhi recruitment and piRNA production, which are enforced by transgenerational cytoplasmic inheritance. (A) Schematic of the tethering experiment. λN-Panx is tethered to a BoxB-containing nascent transcript in the germline of *D. melanogaster* ovaries. The positions of the primers used for ChIP-qPCR are indicated. **(B) Panx tethering triggers Rhi recruitment.** The levels of Rhi, H3K9me3, and HP1 were measured at the reporter by ChIP-qPCR upon tethering of Panx, Wde (a partner of the H3K9 methyltransferase SetDB1), and a control protein (mkate2) in the germline. Both Panx and Wde induce H3K9me3 deposition and HP1 recruitment. However, Rhi is recruited to the reporter upon Panx tethering, but not Wde tethering. Results were measured across three biological replicates and normalized to the Rhi- and H3K9me3-enriched *42AB* region (chr2R: 6323128–6323219). Error bars indicate the standard deviation of the three biological replicates. Statistical significance was estimated by a two-tailed Student’s t-test; n.s. > 0.05, * *p* < 0.05, ** *p* < 0.01, *** *p* < 0.001, **** *p* < 0.0001. **(C) Reporter expression is suppressed upon Panx and Wde tethering.** RT-qPCR was performed upon tethering of Panx, Wde, and the control protein (mkate2), using the primers indicated in panel A. Results were measured across three biological replicates and normalized to *rp49* mRNA levels. Error bars indicate the standard deviation of the three biological replicates. Statistical significance was estimated by a two-tailed Student’s t-test; * *p* < 0.05, *** *p* < 0.001. **(D) Panx tethering converts the reporter into a piRNA-producing locus.** Shown are profiles of piRNAs mapping to the reporter upon Panx and control tethering in the first generation (G1), as well as in flies where tethering was maintained over multiple generations (MG). Total small RNAs as well as oxidation-resistant small RNAs are shown. (Top right) The bar graph shows the levels of reporter piRNAs in each library. Reporter piRNA levels are normalized to total piRNA reads. Error bars indicate the standard deviation of two biological replicates. (Middle) Size distributions of the reporter small RNAs. (Bottom right) Shown is the nucleotide composition of reporter small RNAs within the 23–29 nt piRNA size range. **(E) Upon Panx tethering, Rhi occupancy increases over generations, while H3K9me3 levels remain unchanged.** Rhi and H3K9me3 levels were measured at the reporter using ChIP-qPCR in G1 and multi-generational (MG) Panx tethering experiments. Results were measured across three biological replicates and normalized to the endogenous *42AB* region (chr2R: 6323128–6323219); the *rp49* gene region served as a control. Error bars indicate the standard deviation of the three biological replicates. Statistical significance was estimated by a two-tailed Student’s t-test; n.s. > 0.05, ** *p* < 0.01. **(F) Panx tethering triggers non-canonical transcription of the reporter locus.** RNA FISH was performed using a probe to detect antisense transcripts of the reporter. Scale bar represents 5 μm. **(G) piRNA biogenesis induced by Panx depends on the cytoplasmic inheritance of maternal piRNAs.** (Top) Schematic of the maternal and paternal crosses using parents subjected to multi-generational Panx tethering. (Bottom) The bar graph shows the levels of reporter piRNAs in the progeny of maternal and paternal crosses. Error bars indicate the standard deviation of two biological replicates. **(H) Rhi recruitment depends on the cytoplasmic inheritance of maternal piRNAs.** The levels of Rhi and H3K9me3 were measured at the reporter in the progeny of maternal and paternal crosses using ChIP-qPCR. Results were measured across three biological replicates and normalized to the endogenous *42AB* region (chr2R: 6323128–6323219). Error bars indicate the standard deviation of the three biological replicates. Statistical significance was estimated by a two-tailed Student’s t-test; n.s. > 0.05, ** *p* < 0.01. **(I) Continuous Panx tethering is required to maintain piRNA production and reporter repression.** Following multi-generational Panx tethering, flies were crossed to produce progeny that either retained (Maternal) or lacked (Maternal, no λN-Panx) the tethering machinery. **(Left)** Reporter transcript levels were quantified in the progeny by RT-qPCR and normalized to *rp49* mRNA. Error bars indicate the standard deviation of three biological replicates. **(Right)** Reporter piRNA levels in progeny with or without continued Panx tethering. Error bars indicate the standard deviation of two biological replicates.

To determine whether the deposition of H3K9me3 is, by itself, sufficient for Rhi recruitment, we employed an alternative strategy to install this mark by tethering Windei (Wde), a critical cofactor of the H3K9 methyltransferase dSetDB1^34^. Wde tethering successfully induced transcriptional silencing (Fig. 4C) and triggered both H3K9me3 deposition and HP1 recruitment (Fig. 4B). However, we observed only minimal Rhi enrichment following Wde tethering. These results demonstrate that while H3K9me3 deposition is sufficient to recruit HP1 via a piRNA-independent mechanism, it is insufficient to drive the robust recruitment of Rhi. Instead, Rhi recruitment appears to require the Panx-dependent piRNA pathway rather than the general heterochromatin assembly machinery.

At the chromatin of piCs, Rhi interacts with several partner proteins to initiate non-canonical transcription and the export of piRNA precursors^14^. To determine whether Panx-mediated Rhi recruitment is sufficient to trigger piRNA biogenesis, we profiled ovarian small RNAs. While initial Panx tethering was associated with the emergence of small RNAs from the reporter locus, their levels were relatively low (Fig. 4D). Given that piRNA biogenesis in *Drosophila* is heavily influenced by the cytoplasmic inheritance of maternal piRNAs^11,35,36^, we examined the effects of multi-generational Panx tethering. Remarkably, while H3K9me3 levels remained stable compared to the first generation, multi- generational tethering led to an additional 2.2-fold increase in Rhi occupancy at the reporter (Fig. 4E). Moreover, we detected a significant increase in small RNAs originating from both genomic strands of the reporter and its flanking sequences (Fig. 4D).

Analysis of the nucleotide composition of these reporter-derived small RNAs revealed a strong 1U bias, and their length distribution showed that the majority ranged from 23 to 29 nucleotides—typical of the piRNA size class—with a minor population of 21 nt endo-siRNAs (Fig. 4D). These small RNAs were also resistant to oxidation, indicating they possess the characteristic 3’-terminal 2’-O-methyl (2’-OMe) modification (Fig. 4D). The presence of these *bona fide* piRNAs, coupled with the detection of antisense transcripts by RNA FISH (Fig. 4F), demonstrates that Panx-mediated Rhi recruitment is sufficient to activate non-canonical transcription and transform a naive genomic locus into a functional, dual-strand piC.

Increased Rhi and piRNA levels following multi-generational Panx tethering suggest that the cytoplasmic transmission of piRNA between generations enhances piRNA biogenesis. To further explore the role of maternally inherited piRNAs, we crossed wild-type flies with males and females subjected to multi-generational Panx tethering. While the female offspring of these two crosses are genetically identical, only the progeny of the maternal cross cytoplasmically inherit reporter piRNAs (Fig. 4G). ChIP-qPCR revealed similar levels of the H3K9me3 mark in both progenies; however, the progeny of the maternal cross—those that inherited piRNA—exhibited significantly higher levels of Rhi (Fig. 4H). Furthermore, reporter-derived piRNA levels were 2.1-fold higher in the maternal cross progeny (Fig. 4G). These data demonstrate that the cytoplasmic transmission of piRNA is required to maintain the high levels of Rhi and piRNA biogenesis induced by Panx tethering.

We next investigated whether the high levels of Rhi achieved through multi-generational transmission rendered Panx tethering dispensable. The removal of Panx tethering, even while preserving cytoplasmic piRNA transmission, triggered the loss of piRNA production and derepression of the reporter (Fig. 4I). This indicates that continuous Panx tethering is required to maintain a heterochromatin repression. Overall, our results indicate that while Panx tethering triggers Rhi deposition and subsequent piRNA biogenesis, the efficiency of these processes depends on the continuous cytoplasmic transmission of piRNA across generations.

### Panx recruits Rhi through a dual mechanism of H3K9me3 deposition and direct physical interaction

Our findings suggest that Panx recruits Rhi via a dual mechanism: depositing an H3K9me3 anchor and directly binding Rhi. Although the unstructured N-terminal region of Panx is known to mediate H3K9me3 establishment^25,28,30,31^ (Fig. 5A), how Panx directly recruits Rhi remained unclear. Using co-immunoprecipitation (Co-IP) assays in S2 cells—a system devoid of endogenous germline piRNA components—we demonstrated a physical interaction between Panx and Rhi (Fig. 5B). We further characterized this association using yeast two-hybrid (Y2H) assays, which confirmed a specific physical interaction between Rhi and Panx (Fig. 5A, C). Notably, while both HP1 and Rhi are deposited onto chromatin upon Panx tethering, Y2H assays showed that Panx interacts specifically with Rhi, but not with HP1 (Fig. 5D). Furthermore, no interaction between Panx and Kipf was detected, supporting the model that Kipf operates through a distinct Rhi recruitment pathway (Fig. S3A).

**Figure 5.**
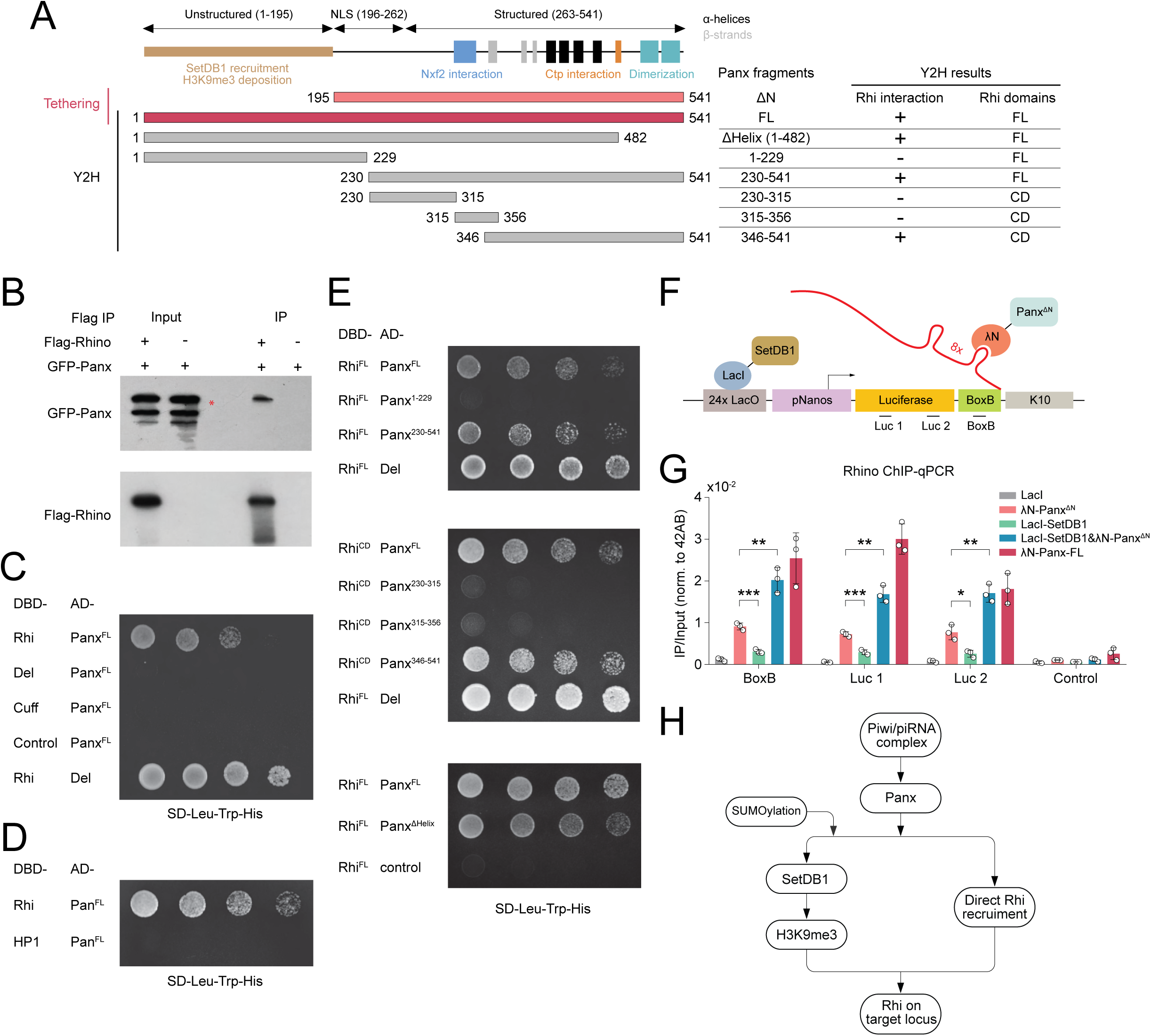
Panx acts as a modular scaffold to coordinate direct Rhi recruitment and chromatin modification. (A) Structure of the Panx protein and dissection of its interaction with Rhi. Schematic of the Panx structure, indicating the unstructured N-terminal region—involved in the recruitment of SetDB1 and establishment of the H3K9me3 mark—as well as other structural elements. The fragments of Panx used in the Y2H and tethering experiments are shown below, along with a summary of the Y2H results. Either full-length (FL) Rhi or the Rhi chromodomain (CD) was used in the Y2H experiments. **(B) Co-immunoprecipitation reveals a Panx-Rhi interaction.** FLAG-Rhi and GFP-Panx were co-expressed in S2 cells, followed by immunoprecipitation using anti-FLAG beads and Western blotting using an anti-GFP antibody. The asterisk indicates the expected size of the GFP-Panx protein. **(C) Panx interacts with Rhi in a Y2H assay.** Y2H experiments were performed to test the interaction between Panx and components of the RDC complex (Rhi, Del, and Cuff). The mkate2 protein served as a control. **(D)** Panx interacts with Rhi but not HP1. **(E) The C-terminal portion of Panx interacts with Rhi.** Y2H results showing the interactions between different fragments of Panx and Rhi. The Panx fragment that binds Rhi (residues 346–541) does not overlap with the N-terminal portion involved in the establishment of the H3K9me3 mark. Panx dimerization is dispensable for the Rhi interaction, as binding is preserved in the Panx^ΔHelix^ truncation. **(F) Schematic of the double-tethering reporter.** Simultaneous DNA and RNA tethering in the *Drosophila* germline was achieved by employing the LacO-LacI (DNA) and λN-BoxB (RNA) systems. DNA tethering was used to tether the H3K9 methyltransferase SetDB1, while RNA tethering was used to tether either full-length or truncated Panx expressed in the germline. **(G) Efficient Rhi recruitment requires both direct Panx-Rhi interaction and H3K9me3 deposition.** Rhi occupancy was measured at the reporter by ChIP-qPCR following single or double tethering of indicated proteins. Error bars indicate the standard deviation of three biological replicates. Statistical significance was estimated by a two-tailed Student’s t-test; n.s., not significant, * p < 0.05, ** p < 0.01, *** p < 0.001. **(H) Two-pronged model of piRNA-guided Rhi recruitment.** Panx is tethered to nascent transcripts following their recognition by the nuclear Piwi/piRNA complex. The N-terminal portion of Panx mediates the establishment of the H3K9me3 mark for Rhi binding through the SUMO-dependent recruitment of the H3K9 methyltransferase SetDB1. The C-terminal portion of Panx interacts directly with Rhi, mediating its deposition onto H3K9me3-enriched chromatin.

To define the structural basis of Panx-Rhi interaction, we mapped the domains of Panx responsible for Rhi binding. Y2H assays using truncated fragments revealed that the interaction is mediated by the Panx C-terminal region (residues 346–541; Fig. 5A and 5E). While this region includes alpha-helices required for Panx dimerization, their deletion did not disrupt Rhi binding. On the Rhi side, we mapped the interaction with Panx to the chromodomain (Fig. S3B) and found that disrupting its dimerization capacity^11^ abolished the interaction (Fig. S3C). Using AlphaFold-Multimer, we generated five candidate structural models of the Rhi-chromodomain-dimer/Panx complex. Mutational analysis in Y2H assays validated a single conformation (Model 3): while mutations targeting interfaces in the other models had no effect, substituting seven specific contact residues from Model 3 with alanines disrupted the interaction (Fig. S3D, E). Taken together, our experiments reveal that Panx binds Rhi via its C-terminal portion, while the function of Panx in establishing the H3K9me3 mark has been previously mapped to its N-terminal region (residues 1–195)^25,28,30,31^. Thus, Panx utilizes two distinct, modular regions to coordinate Rhi recruitment: a disordered N-terminus for chromatin modification and a structured C-terminus for direct Rhi binding (Fig. 5E).

Mapping the two functions of Panx—Rhi interaction and heterochromatin establishment—to distinct regions allows for a detailed exploration of their respective roles in Rhi recruitment. To investigate the necessity of these regions, we used a modified tethering reporter which allows DNA and RNA tether to the reporter simultaneously (Fig. 5F). As expected, the N-terminal truncation of Panx largely abolished H3K9me3 installation (Fig. S3F). However, this truncated protein—which retains the C-terminal region required for Rhi interaction—still recruited Rhi to the reporter, albeit at reduced levels. This indicates that the Panx-Rhi interaction mediated by the C-terminal region can drive moderate Rhi recruitment even in the absence of a robust chromatin anchor (Fig. 5G). Conversely, tethering the methyltransferase SetDB1 to install the H3K9me3 mark resulted in significantly less Rhi recruitment than tethering the truncated Panx.

Remarkably, the simultaneous tethering of truncated Panx and SetDB1 restored Rhi recruitment to levels comparable to those of full-length Panx (Fig. 5G). This demonstrates that the two functions of Panx—chromatin mark installation and direct Rhi recruitment—are physically separable, and that the former can be replaced by general heterochromatin machinery. Collectively, these experiments suggest that the piRNA-guided Piwi/Panx complex directs Rhi to its targets with a specificity and efficiency that far exceeds that provided by the H3K9me3 mark alone (Fig. 5H).

## Discussion

### Beyond the histone code: RNA-directed recruitment as a paradigm for reader specificity

The traditional understanding of chromatin regulation posits a simple “lock-and-key” model, where diverse histone modifications serve as precise docking sites for their cognate reader proteins. While this model holds true for some factors—such as HP1, whose genomic profile closely matches the ubiquitous distribution of H3K9me3—it fails to capture the complexity of other chromatin readers, such as the HP1 paralog Rhi. As extensive genomic investigations across various organisms have shown, readers are frequently excluded from regions where their target marks are abundant^37,38^. This genomic discordance indicates that the presence of a histone mark is often necessary but insufficient for recruitment, requiring additional regulatory layers to define binding specificity. Our findings demonstrate that Rhi specificity is not an intrinsic property of its interaction with the H3K9me3 mark, but is instead imposed by sequence-specific, small RNA-directed targeting.

Several distinct mechanisms have been identified that explain how reader specificity is restricted *in vivo*. For instance, the reader BPTF requires the DNA-binding pioneer factor FOXA1 to first open compact chromatin before it can access its target marks, illustrating a hierarchical gating mechanism^39^. Other readers rely on combinatorial signals, such as UHRF1, which requires reciprocal cooperativity between H3K9me3 and hemimethylated DNA to achieve stable binding^40,41^. Additionally, internal protein structures can dictate occupancy; the *Arabidopsis* reader EBS utilizes an auto-inhibition loop to completely decouple its binding potential from the physical distribution of its target marks^42^.

Most relevant to our findings, non-coding RNAs have emerged as critical guides for overriding the default histone landscape. For example, the lncRNA *Khps1* forms an RNA:DNA:DNA triplex to sequence-specifically recruit the p300 reader to a promoter, independent of pre-existing acetylation marks^43^. Our study establishes the piRNA-directed recruitment of Rhi as a powerful new paradigm within this RNA-guided class. While H3K9me3 provides a foundational platform, the true specificity for Rhi is driven either by DNA-binding factors like Kipferl^10^ or by piRNAs that recognize nascent transcripts and physically tether Panx to the locus. This demonstrates that small RNAs can effectively upgrade a ubiquitous histone code to dictate the precise genomic distribution of a chromatin reader.

The observation that Panx, Kipf, and the H3K27 methyltransferase E(z) compete for Rhi recruitment demonstrates that the partitioning of this reader across the genome is a tightly contested process. The redistribution of Rhi to Kipf-dependent sites upon Panx depletion (Fig. 2) indicates that the total pool of Rhi protein is a limited resource. This competition likely reflects an evolutionary strategy to counter diverse genomic threats by maintaining multiple recruitment pathways to ensure a flexible defense network.

### The Panx as a modular scaffold

To execute RNA-guided specificity of Rhi recruitment, the piRNA pathway relies on the unique structural properties of Panx, which acts as a modular, two-pronged scaffold. Its N-terminus is responsible for recruiting the methyltransferase machinery to install the H3K9me3 anchor^25,28,30,31^, while its C-terminus simultaneously engages in the direct physical recruitment of Rhi (Fig. 5). This dual-mode mechanism ensures that the installation of the mark and the docking of the reader are perfectly coordinated and occur exclusively at genomic regions recognized by piRNAs (Fig. 5H).

While the direct Panx-Rhi interaction can drive moderate recruitment—and H3K9me3 alone is insufficient—the simultaneous presence of both restores robust Rhi occupancy (Fig. 5G). This mirrors the “cooperative logic” observed in other specific epigenetic regulators, such as UHRF1, which requires both H3K9me3 and hemimethylated DNA to achieve stable chromatin interaction^44^. We propose a “synergy model” for Rhi recruitment: H3K9me3 provides the necessary avidity (a broad, stable platform), while the direct Panx-Rhi physical bridge provides the sequence specificity (Fig. 5). Without the Panx, Rhi cannot distinguish piCs from general heterochromatin; without the H3K9me3 platform, the recruitment may be too transient to sustain robust occupancy.

*In vivo*, Panx-dependent Rhi recruitment may be further enhanced by biomolecular condensation. The C-terminal region of Panx—the exact domain required for Rhi interaction—has been shown to facilitate liquid-liquid phase separation (LLPS) in the presence of RNA *in vitro*^28^. Consistent with this, Rhi and its partners form distinct, concentrated foci within the nucleus, indicative of condensate formation^7,45^. Panx-driven condensates likely create a specialized microenvironment that favors the cooperative recruitment and retention of Rhi, ensuring that piRNA clusters assemble efficiently and remain distinct from generic, H3K9me3-enriched heterochromatin.

### Transgenerational epigenetic memory via a self-reinforcing piRNA loop

The ultimate biological consequence of targeted Rhi recruitment is the establishment of a robust, transgenerational epigenetic memory that drives piRNA biogenesis. As demonstrated by our tethering assay, Panx recruitment is sufficient to trigger Rhi deposition and establish a *de novo* piRNA clusters (Fig. 4). However, this initial establishment is dramatically amplified by the transgenerational inheritance of maternal piRNAs. Once localized, Rhi drives the non-canonical transcription of piRNA precursors, feeding the biogenesis machinery to produce mature piRNAs. Crucially, because these newly minted piRNAs are maternally inherited by the progeny^35^, they endow the developing embryo with a pre-programmed set of sequence-specific guides. These inherited piRNAs then guide the Piwi-Panx complex back to the exact same genomic loci to initiate H3K9me3 deposition and Rhi recruitment anew. This creates a self-reinforcing feed-forward loop: piRNAs guide Rhi deposition, and Rhi deposition produces more piRNAs. Through this mechanism, the piRNA pathway bypasses the limitations of the histone code, utilizing RNA not merely as a targeting guide for a chromatin reader, but as an epigenetic memory molecule that guarantees the continuous defense of the genome across generations.

## Acknowledgements

We thank Katalin Fejes Tóth and members of the Aravin and Fejes Toth labs for discussion and comments. We thank Julius Brennecke for providing the Rhino antibody. We thank Igor Antoshechkin (Caltech) for helping with sequencing. This work was supported by grants from National Institutes of Health (R01 GM097363) and by the HHMI Faculty Scholar Award to A.A.A and the Sandler Program for Breakthrough Biomedical Research to P.H.

## Materials and Methods

### Drosophila stocks

All fly lines used in this study are listed in Supplementary Table 1. Flies were maintained on standard medium at 25°C under conventional laboratory conditions.

### Transgenic flies

To generate the pUbiquitin-GFP-NLS-SV40 construct, the Ubiquitin promoter, GFP-NLS, and SV40 polyadenylation signal were PCR-amplified and assembled into the EcoRI- and BamHI-digested pBS-KS-attB1-2 vector using Gibson Assembly. The resulting construct was integrated into genomic loci at chr2R:6338399 (BDSC #43121) and chr2R:6439020 (BDSC #44828).

For λN tethering constructs (pUAS-λN-Flag-Cuff, -Piwi, -mkate2, -Panx, -Wde, and pNanos-λN-Flag-Panx, -mkate2), corresponding cDNAs were cloned into pENTR™/DTOPO® entry vectors (Invitrogen) and recombined into Gateway-compatible destination vectors containing ΦC31 attB sites, mini-white markers, and either UASp or nanos promoters fused to an N-terminal λN-Flag cassette.

To generate the LacO-pNanos-luciferase-BoxB-K10 reporter, the nanos promoter, luciferase coding sequence, and eight BoxB repeats were assembled into a ΦC31-compatible backbone using Gibson Assembly. A 24× LacO array was subsequently inserted by partial XbaI digestion of pSV2-DHFR-8.32 provided by A.S. Belmont^46^ and ligation into the reporter vector. The final construct was integrated at the 76A2 landing site (VK00013, BDSC #9732).

Unless otherwise specified, all constructs were inserted into the attP40 (25C6) or attP2 (68A4) landing sites. Transgenesis was performed by BestGene. A complete list of transgenic lines is provided in Supplementary Table 1.

### RNA HCR-FISH

RNA fluorescence in situ hybridization using hybridization chain reaction (HCR-FISH) was performed with modifications from previously described protocols^47,48^. Probes were designed and synthesized by Molecular Technologies. Fluorophores Alexa Fluor 546, 594, and 647 were used for detection. Images were acquired using a ZEISS LSM880 confocal microscope and analyzed with ZEN software.

### Immunofluorescence microscopy

Immunofluorescence was conducted following the previously described protocol^15,49^. Ovaries were fixed in 4% paraformaldehyde (PFA) in PBS supplemented with 0.1% Triton X-100 (PBST) for 20 min. Following fixation, samples were blocked in PBST containing 1% BSA for 1 h at room temperature and then incubated with primary antibodies overnight at 4°C. Primary antibodies used were anti-H3K9me3 (Abcam, ab8898) and anti-HP1a (DSHB, C1A9). After primary antibody incubation, samples were washed in PBST and incubated with Alexa Fluor–conjugated secondary antibodies (Alexa Fluor 546 or 647) at a 1:1000 dilution for 3 h at 4°C. Samples were subsequently washed and stained with 1 μM DAPI (Sigma-Aldrich). Finally, ovaries were washed in PBS and mounted in VECTASHIELD antifade mounting medium for imaging. Imaging was performed using a ZEISS LSM880 confocal microscope.

### ChIP-seq and ChIP-qPCR

All ChIP experiments were conducted according to a previously established protocol^13^, Antibodies used include anti-H3K9me3 (Abcam, ab8898), anti-HP1a (DSHB, C1A9), and anti-Rhino (gift from the Brennecke laboratory). For ChIP-qPCR, enrichment was quantified using SYBR Green-based qPCR (MyTaq HS Mix, BioLine) on a Mastercycler® ep Realplex system (Eppendorf). Ct values were calculated from technical duplicates. ChIP signals were normalized to input and to a region in 42AB cluster (chr2R: 6,323,128 – 6,323,219). Primer sequences are listed in Supplementary Table 2.

### Small RNA-seq

Small RNA cloning and sequence experiments were followed the previous protocol^35^. Total RNA was extracted from dissected ovaries using TRIzol (ThermoFisher, #15596018). 4µg total RNA were size-selected on a 15% polyacrylamide gel, and RNA fragments between 19–29 nt were isolated. For oxidation assays (Fig. 4D), RNA samples were treated with 200 mM sodium periodate at 25°C for 30 minutes prior to library preparation. Libraries were constructed using the NEBNext Small RNA Library Prep Kit (NEB, E7330S) and sequenced on an Illumina HiSeq 2500 platform.

### RT-qPCR

Total RNA was extracted from approximately 20 dissected ovaries using TRIzol TRIzol (ThermoFisher #15596018). 1µg of RNA was treated with DNase I and reverse transcribed using SuperScript III (Invitrogen, #18080044). Quantitative PCR was performed using MyTaq HS Mix (BioLine, BIO-25046) with SYBR Green detection. Ct values were derived from technical duplicates, and expression levels were normalized to *rp49* mRNA. Primer sequences are listed in Supplementary Table 2.

### Co-immunoprecipitation

Co-immunoprecipitation protocol was modified from previous study^15^. Cells were co-transfected with pActin-Flag-Rhi and pActin-GFP-Panx constructs, with pActin-GFP-Panx alone serving as a control. Protein complexes were immunoprecipitated using anti-Flag M2 beads (Sigma, M8823-5ML). Western blot analysis was performed using anti-Flag (1:2000) and anti-GFP (1:1000) antibodies in 5% non-fat milk.

### Yeast two-hybrid assay

The yeast competent cells (AH109) were prepared following the previous protocol^50^. Protein–protein interactions were assessed using yeast two-hybrid (Y2H) assays. Coding sequences were cloned into pGADCg (activation domain) and pGBKCg (DNA-binding domain) vectors. Constructs were co-transformed into AH109 yeast cells and plated on SD-Leu-Trp selection medium. Colonies were cultured to OD600 ≈ 3, serially diluted (1:3, 1:9, 1:27), and plated onto SD-Leu-Trp-His plates to assess interaction-dependent growth.

### Modeling of the Rhi-Panx interaction

Structural predictions of the Rhi-Panx complex were generated using LocalColabFold (https://github.com/YoshitakaMo/localcolabfold) ^51^. The complex was modeled using a 2:1 stoichiometry of the Rhi chromodomain to Panx. Five candidate structural models were generated, from which predicted interface residues were identified and selected for targeted mutagenesis to be validated in subsequent yeast two-hybrid (Y2H) assays.

### Bioinformatic analysis of ChIP-seq and small RNA libraries

ChIP-seq reads were processed following previously described pipelines^35^. Adapter trimming and quality filtering were performed using Trimmomatic (version 0.33)^52^ and cutadapt (version 1.15)^53^. Reads were mapped to the *Drosophila melanogaster* dm6 genome and relevant vector sequences using Bowtie (v1.0.1)^54^ (parameters: -v 2 -k 1 -m 1 -t --best -y --strata). Mitochondrial reads were excluded. Coverage tracks and enrichment profiles were generated using customized scripts and deepTools^55^. Regions blacklisted by ENCODE^56^ were excluded from enrichment analysis. Read counts were computed in fixed genomic bins using deepTools2 and BEDOPS^57^, and visualization was performed in MATLAB. All scripts employed in this analysis can be found on GitHub, https://github.com/Peng-He-Lab/Luo_2025_piRNA For small RNA-seq analysis were followed the previous protocol^35^. Small RNA reads were trimmed and filtered to remove reads shorter than 20 nt. Subsequently, reads of specific lengths were extracted: 21-22nt (siRNA), 23-29 nt (piRNA), and 21-30 nt (small RNA). Reads were mapped to the dm6 genome using Bowtie (parameters: -v 0 -a -m 1 - t --best --strata). After removal of mitochondrial reads, read densities were calculated using deepTools2 and BEDOPS. The ping-pong signature was inferred using previously published method^58^. Genome coverage and insertion-specific analyses were generated using MATLAB. The mapping scripts are available on GitHub, https://github.com/Peng-He-Lab/Luo_2025_piRNA/blob/main/ChIP-seq.md

### Identification of *de novo* TE insertions

*De novo* TE insertions were identified from ChIP-seq input libraries of *Panx* and *white* GLKD using a custom pipeline that captures chimeric reads spanning TE-genome junctions. After adapter trimming and quality filtering, the first 20 nt of each read were extracted and aligned to a *D. melanogaster* TE consensus index using Bowtie (v1.0.1) (parameters: -v 2 -k 1); reads whose 5’ ends mapped to a TE consensus were retained, and their full-length sequences were recovered. To exclude reads derived entirely from internal TE sequences, the last 20 nt of these full-length reads were aligned back to the TE consensus index, and only reads whose 3’ ends failed to map (parameter: --un) were carried forward. The unmapped 3’ fragments were then aligned to the dm6 genome (parameters: -v 0 -k 1 -m 1) to identify uniquely mapping insertion sites. To exclude reads originating from endogenous, full-length TE copies present in the reference genome, the corresponding full-length reads were realigned to dm6, allowing up to three mismatches (parameter: -v 3); aligned reads were discarded, and only reads that failed to map were retained (parameter: --un). The 3’ ends of these remaining reads were then remapped to the genome (parameters: -v 0 -k 1) to define precise insertion coordinates. Genomic coordinates were converted to BED format, and overlapping intervals were merged using bedtools (v2.30.0). To restrict the analysis to euchromatic regions and to remove signal from pre-existing repetitive elements, intervals overlapping the dm6 RepeatMasker track (downloaded from the UCSC Genome Browser) were excluded with bedtools intersect -v. This pipeline yielded 287 *de novo* TE insertions across the *Panx* and *white* GLKD samples.

## Data availability

All sequencing data generated in this study are deposited in GEO under accession number GSE297619. Previously published datasets used in this study include HP1 ChIP-seq GSE115277^59^, Kipf and Rhi ChIP-seq GSE202468^10^, and H3K27me3 ChIP-seq GSE247156^32^.

## Author Contributions

Y.L. and A.A.A. designed the experiments; Y.L. performed all experiments; W. Z. and P.H. analyzed the *de novo* TE insertion; Z. J. performed the *in vitro* protein interaction assays; B. G. verified the Y2H results; X. H. performed the Co-IP experiments; H. W. performed the Panx-Rhi complex structure prediction by AlphaFold2; Y. H. designed the *in vitro* protein interaction assays; Y.L. and P.H. analyzed the data; Y.L. and A.A.A. wrote the paper.

---

**Figure S1.**
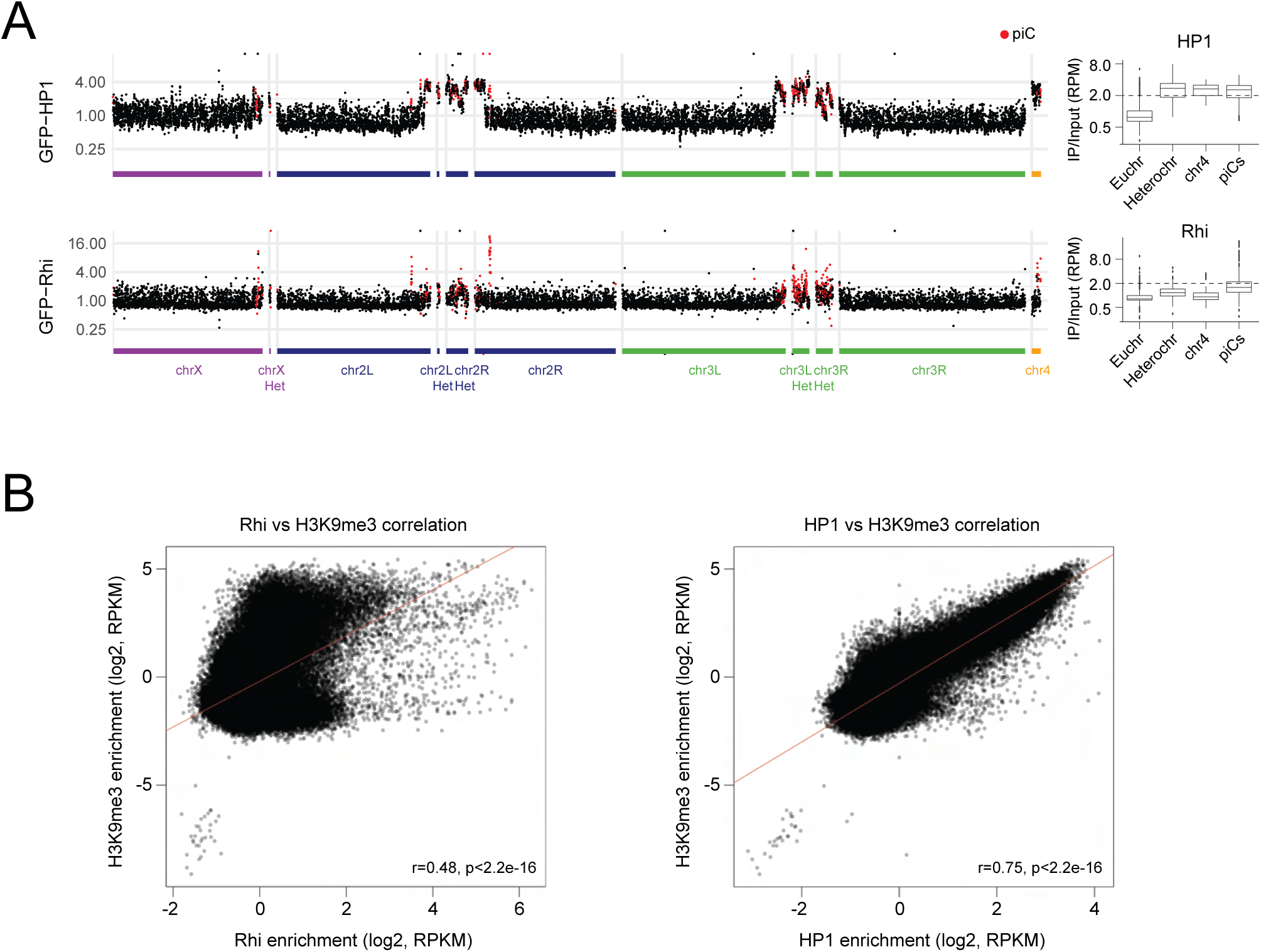
related to Figure 1. (A) Rhi and HP1 show distinct chromatin profiles in germline. ChIP-seq profiling of GFP-tagged Rhi and HP1 was performed using anti-GFP antibodies on ovaries dissected from flies expressing *UASp-GFP-Rhi* and *UASp-GFP-HP1* transgenes driven in the germline by the maternal *tubulin-GAL4* driver. Data points correspond to input-normalized ChIP signals in 10 kb genomic intervals. Intervals overlapping piCs are marked in red. Boxplots show the distributions of ChIP signals measured in 10 kb genomic intervals within euchromatin (Euchr), pericentric heterochromatin (Heterochr), heterochromatinized chromosome 4 (chr4) and piRNA clusters (piCs). Results were averaged from two biological replicates. **(B) Genome-wide correlation of H3K9me3 with Rhi and HP1.** Scatter plots comparing the levels of H3K9me3 versus Rhi, and H3K9me3 versus HP1, across 1-kb genomic intervals. Data points represent the average input-normalized ChIP-seq signal (RPKM) from two biological replicates.

**Figure S2.**
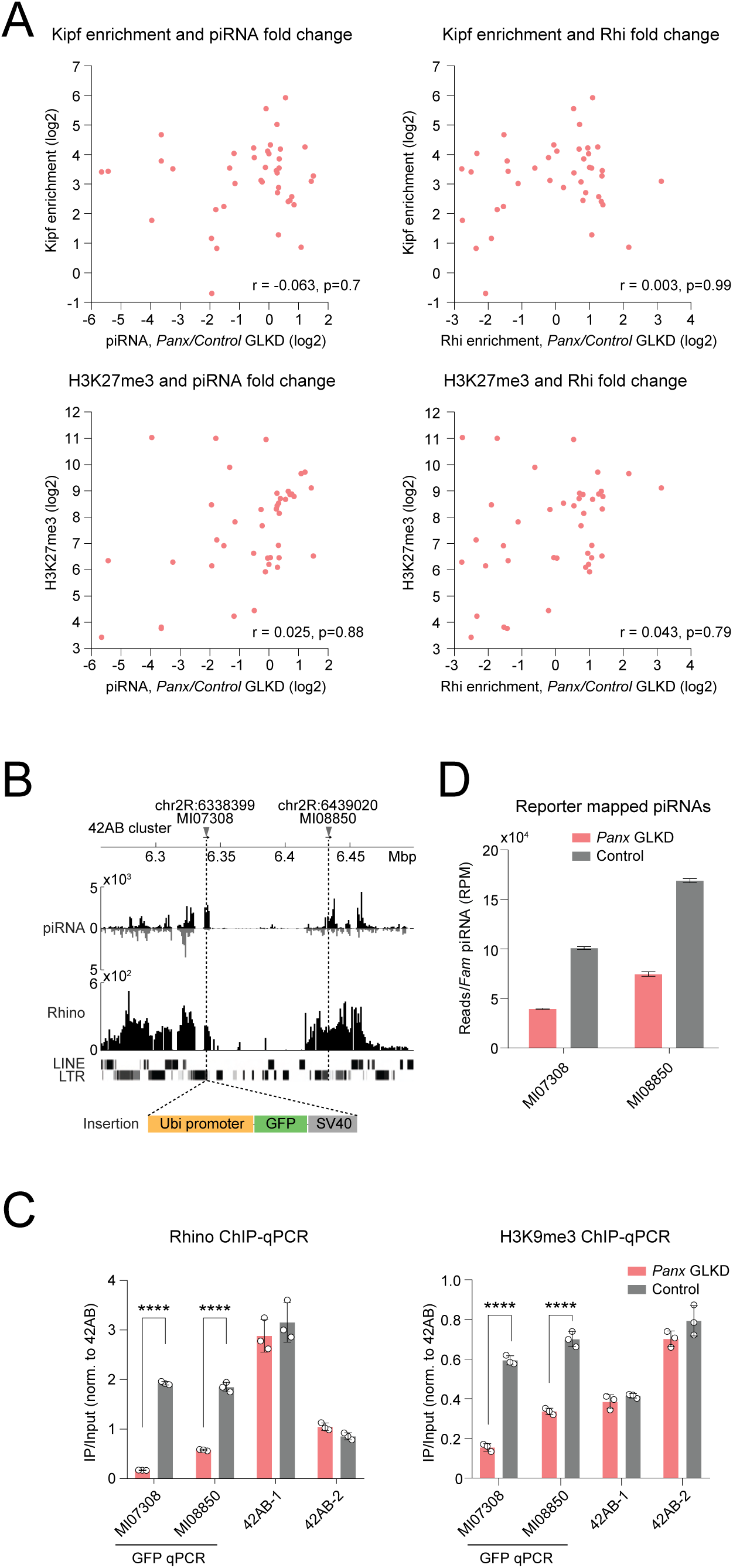
related to Figure 2. Depletion of Panx reduces Rhi occupancy and piRNA production from transgenes integrated into *42AB* cluster. **(A) Occupancy of Kipf or H3K27me3 does not predict piC sensitivity to *Panx* GLKD.** Scatter plots comparing baseline levels of Kipf or H3K27me3 on individual piCs to the changes in piRNA abundance and Rhi occupancy following *Panx* GLKD. Data points represent the average input-normalized signal (RPKM) from two biological replicates. **(B) *Ubiquitin-GFP* transgenes integrated into *42AB* cluster.** *Ubiquitin-GFP* constructs were introduced at two sites within the *42AB* cluster using the Minos-mediated integration cassette (MiMIC) system. **(C) *Panx* is required for Rhi occupancy and H3K9me3 at transgenes integrated into the *42AB* cluster.** ChIP-qPCR analysis of Rhi and H3K9me3 levels at the transgenes upon *Panx* GLKD. Results were measured across three biological replicates and normalized to the endogenous *42AB* region (chr2R: 6449408–6449518). Error bars indicate the standard deviation of three biological replicates. Statistical significance was estimated by a two-tailed Student’s t-test; **** *p* < 0.0001. |**(D)** *Panx* is essential for piRNA biogenesis at transgenes integrated into the *42AB* **cluster.** The bar graph shows piRNA levels from the transgenes upon *Panx* GLKD. piRNA levels are normalized to *Flam* piRNA reads. Error bars indicate the standard deviation of two biological replicates.

**Figure S3.**
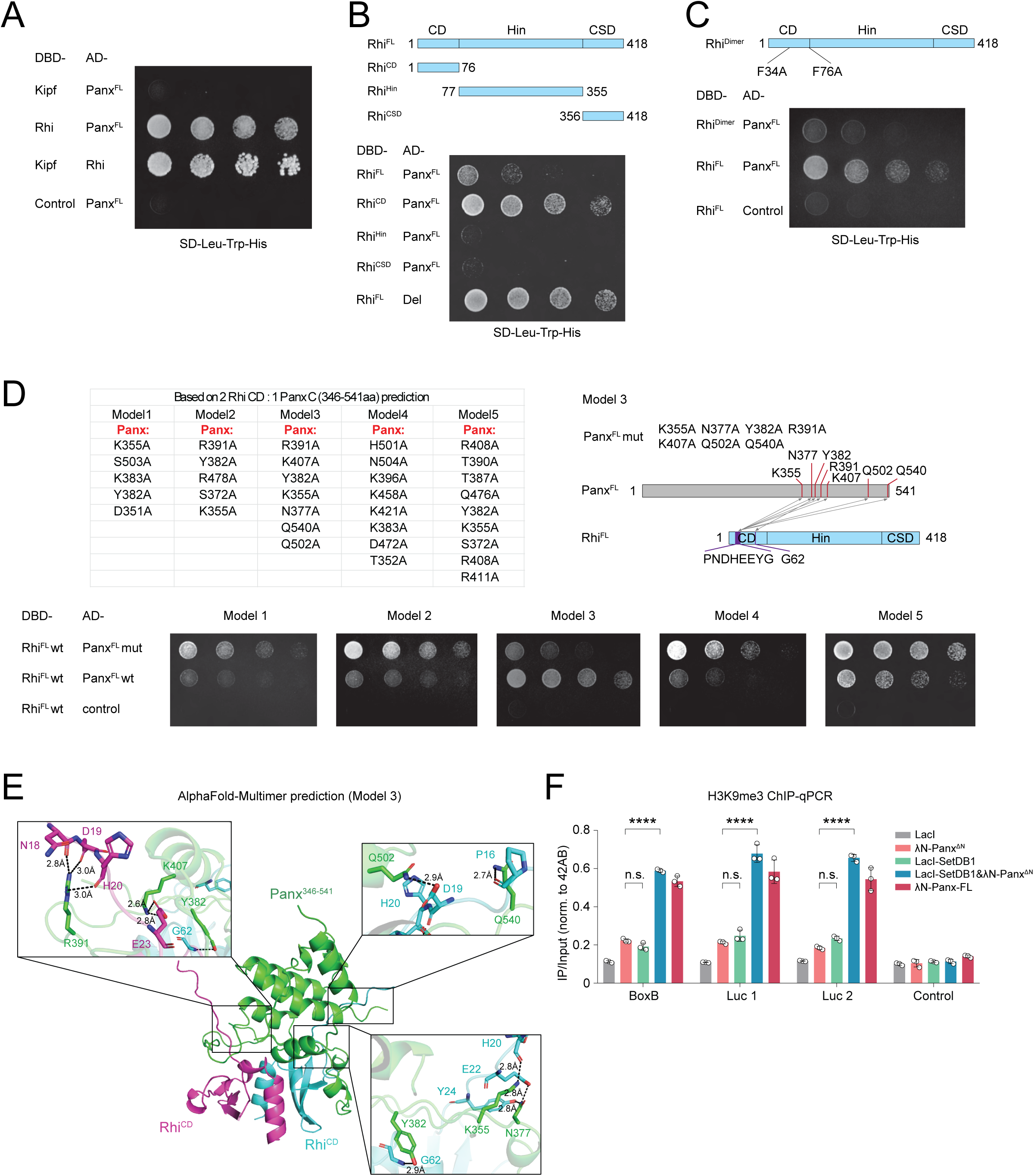
Related to Figure 5. **(A) Panx does not interact with Kipf in a Y2H assay.** The mkate2 protein served as a control. **(B) The Rhi chromodomain interacts with Panx.** Y2H experiments were performed using full-length Rhi, the chromodomain (CD), the hinge domain (Hin), and the chromoshadow domain (CSD). The Rhi-Del interaction was used as a positive control. **(C) Rhi dimerization is required for the Panx-Rhi interaction.** Mutations that disrupt the dimerization of Rhi chromodomain (F34A, F76A) abolish its interaction with Panx. The mkate2 protein served as a control. **(D) Validation of the AlphaFold-predicted Panx-Rhi interface.** Yeast two-hybrid (Y2H) mutational analysis of the Panx-Rhi interaction testing five candidate AlphaFold-Multimer structural models. Only the mutation set designed to disrupt the predicted interface in Model 3 successfully abrogates Panx-Rhi binding. mKate2 served as a negative control. **(E) Structural model of the Panx-Rhi complex.** AlphaFold-Multimer prediction (Model 3) of the interaction between the Panx central region (residues 346–541, green) and two Rhi chromodomains (magenta and cyan). Unstructured residues (pLDDT score < 20) are omitted for clarity. Insets highlight predicted polar contacts at the binding interfaces. **(F) Establishment of the H3K9me3 mark in double-tethering experiments.** H3K9me3 levels were measured at the reporter by ChIP-qPCR following single or double tethering of the indicated proteins in the experiments shown in Figure 5G. Error bars indicate the standard deviation of three biological replicates. Statistical significance was estimated by a two-tailed Student’s t-test; n.s., not significant (*p* > 0.05), **** *p* < 0.0001.

**Table S1.**
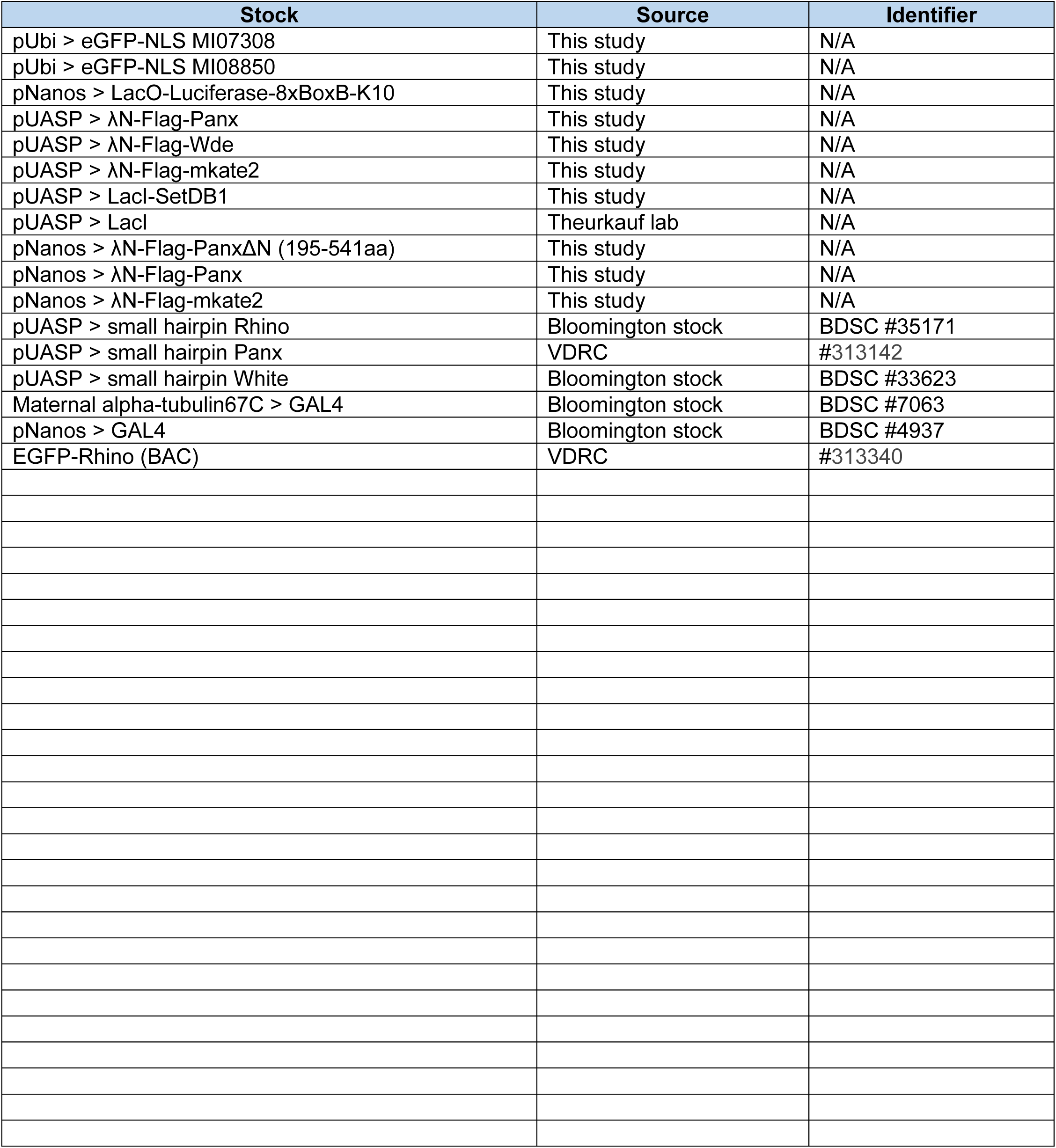
Drosophila melanogaster stocks.

**Table S2.**
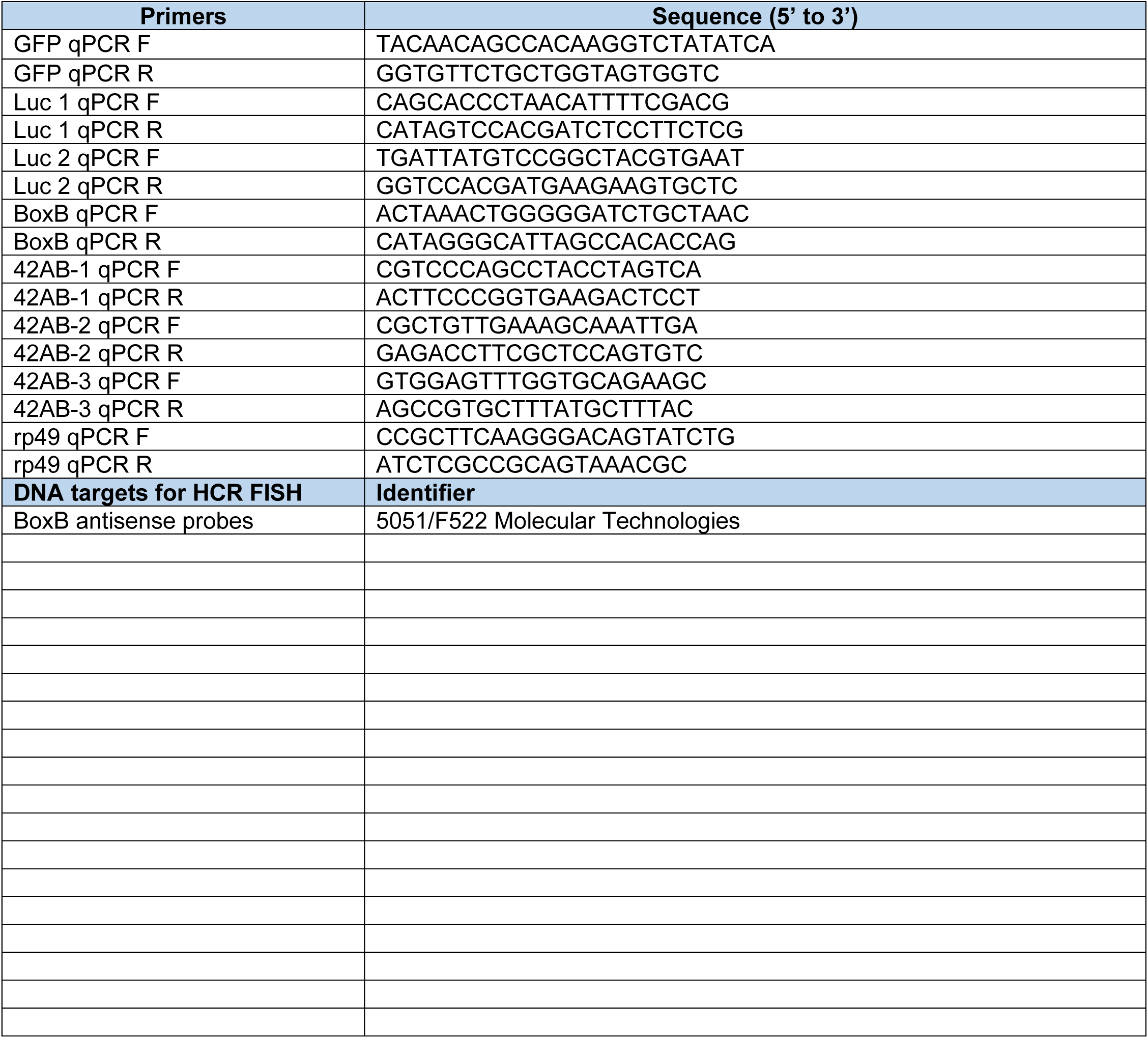
Primers.

